# A Strategy to Treat COVID-19 Disease with Targeted Delivery of Inhalable Liposomal Hydroxychloroquine: A Non-clinical Pharmacokinetic Study

**DOI:** 10.1101/2020.07.09.196618

**Authors:** Tien-Tzu Tai, Tzung-Ju Wu, Huey-Dong Wu, Yi-Chen Tsai, Hui-Ting Wang, An-Min Wang, Sheue-Fang Shih, Yee-Chun Chen

## Abstract

Severe acute respiratory syndrome coronavirus 2 (SARS-CoV-2) is a newly identified pathogen causing coronavirus disease 2019 (COVID-19) pandemic. Hydroxychloroquine (HCQ), an antimalarial and anti-inflammatory drug, has been shown to inhibit SARS-CoV-2 infection *in vitro* and tested in clinical studies. However, lung concentration (6.7 µg/mL) to predict the *in vivo* antiviral efficacy might not be achievable with the currently proposed oral dosing regimen. Further, a high cumulative doses of HCQ may raise concerns of systemic toxicity, including cardiotoxicity. Here, we described a non-clinical study to investigate the pharmacokinetics of a novel formulation of liposomal HCQ administrated by intratracheal (IT) instillation in Sprague-Dawley (SD) rats which achieved 129.4 µg/g (C_max_) in the lung. Compared to unformulated HCQ administered intravenous (IV), liposomal HCQ with normalized dose showed higher (∼30-fold) lung exposure, longer (∼2.5-fold) half-life in lung, but lower blood exposure with ∼20% of C_max_ and 74% of AUC and lower heart exposure with 24% of C_max_ and 58% of AUC. In conclusion, the pharmacokinetics results in an animal model demonstrate the proof of concept that inhalable liposomal HCQ may provide clinical benefit and serve as a potential treatment for COVID-19.

## INTRODUCTION

Hydroxychloroquine (HCQ), an antimalarial and anti-inflammatory drug, is inexpensive, safe, and well tolerated by most patient populations, including those with chronic diseases or immunocompromised status. HCQ is a weak diprotic base that can pass through the lipid cell membrane and preferentially concentrate in acidic cytoplasmic vesicles. HCQ is being studied to prevent and treat coronavirus disease 2019 (COVID-19), possibly via blocking the interactions between virus and angiotensin-converting enzyme-2 (ACE-2) receptor as well as sialic acids receptor and has shown potential *in vitro* (1, 2) and preliminary clinical results (3, 4). As of Jul 8, 2020, a total of 232 studies involves HCQ use among 2478 clinical trials for COVID-19 registered with the clinicaltrials.gov.

However, the effective *in vivo* levels as well as the optimal dosing regimen of HCQ for treating COVID-19 remains unclear. The *in vitro* EC_50_ values (0.72 to 17.31 µM) proposed for optimized dosing regimens are based on extracellular drug concentration (1, 2). A higher lung (intracellular) concentration to predict the *in vivo* antiviral efficacy was suggested (5). The HCQ concentration (6.7 µg/mL) required to clear 100% of SARS-CoV-2 *in vitro* might not be achievable with the currently proposed oral dosing regimen of 800 mg HCQ sulfate orally daily, followed by a maintenance dose of 400 mg given daily for 4 days (2, 6). Further, a high cumulative doses of HCQ may raise concerns of systemic toxicity, including cardiotoxicity (7).

An alternative strategy is needed to bridge the gap between *in vitro* and clinical use of a potentially effective agent in the context of COVID-19 pandemic. A drug delivery directly to the respiratory tracts while minimizing the systemic exposure is a desirable alternative (5). In this report, a proof-of-concept study was conducted to evaluate the systemic pharmacokinetics (PK) profiles and tissue distribution following a single dose of inhalable liposomal HCQ in a rat model and compared to that of unformulated HCQ through intravenous (IV) delivery.

## RESULTS

### HCQ pharmacokinetics in lung

Upon IT administration of liposomal HCQ, significantly more HCQ disposition (Fig. 1A) and longer half-life in lung was observed compared to that those of HCQ by either IV or IT administration (37.5 hours vs. 15.2 hours and 17.7 hours for HCQ-IV and HCQ-IT, respectively) (Table 1). Notably, even at 72 hours post-dose, liposomal HCQ-IT had 77-fold (to HCQ-IV) and 43-fold (to HCQ-IT) higher HCQ concentrations in lung. A single dose of 0.284 mg liposomal HCQ-IT achieved 129.4 μg/g in C_max_ and 4193.2 h*μg/g in AUC_0-72_ and showed overall greater lung exposure with 29-fold in C_max_ and 35-fold in AUC_0-72_ compared to HCQ-IV with normalized dose (Table 2).

**TABLE 1.** Pharmacokinetic parameters of HCQ in rat lung, blood and heart after a single administration of HCQ-IV, HCQ-IT or liposomal HCQ-IT.

| PK parameters | Lung |  |  | Blood<sup>f</sup> |  |  | Heart |  |  |
| --- | --- | --- | --- | --- | --- | --- | --- | --- | --- |
|  | HCQ-IV | HCQ-IT | Liposomal HCQ-IT | HCQ-IV | HCQ-IT | Liposomal HCQ-IT | HCQ-IV | HCQ-IT | Liposomal HCQ-IT |
| T<sub>max</sub> (h)<sup>a</sup> | 0.25 | 0.25 | 0.25 | 0.25 | 0.25 | 0.25 | 0.25 | 0.25 | 24 |
| C<sub>max</sub><br>(μg/g)/(ng/mL)<sup>b</sup> | 9.4 | 47.8 | 129.4 | 433.4 ± 63.6 | 476.7 ± 63.2 | 42.5 ± 18.5 | 4.5 | 3.8 | 0.5 |
| AUC<sub>0-t</sub><sup>c</sup><br>(h*μg/g)/(h*ng/mL)<sup>d</sup> | 251.6 | 328.6 | 4193.2 | 2333.2 ± 247.6 | 2257 ± 420.7 | 827.6 ± 286.0 | 67.2 | 64.2 | 32.1 |
| AUC<sub>∞</sub><br>(h*μg/g)/(h*ng/mL)<sup>e</sup> | 259.9 | 345.8 | 5760.7 | 2434.8 ± 291.4 | 2538.6 ± 509.6 | 1234.1 ± 323.6 | - | - | - |
| t<sub>1/2</sub> (h) | 15.2 | 17.7 | 37.5 | 15.3 ± 2.1 | 22.1 ± 5.7 | 55.2 ± 13.0 | - | - | - |

<sup>a</sup>T<sub>max</sub> was presented as median for blood
<sup>b</sup>C<sub>max</sub> unit: μg/g for lung and heart; ng/mL for blood
<sup>c</sup>AUC<sub>0-72</sub> for blood and lung; AUC<sub>0-24</sub> for heart
<sup>d</sup>AUC<sub>0-t</sub> unit: h\*μg/g for lung and heart; h\*ng/mL for blood
<sup>e</sup>AUC<sub>∞</sub> unit: h\*μg/g for lung and heart; h\*ng/mL for blood
<sup>f</sup>Pharmacokinetic parameters was presented as mean ± SD for blood

**TABLE 2.**
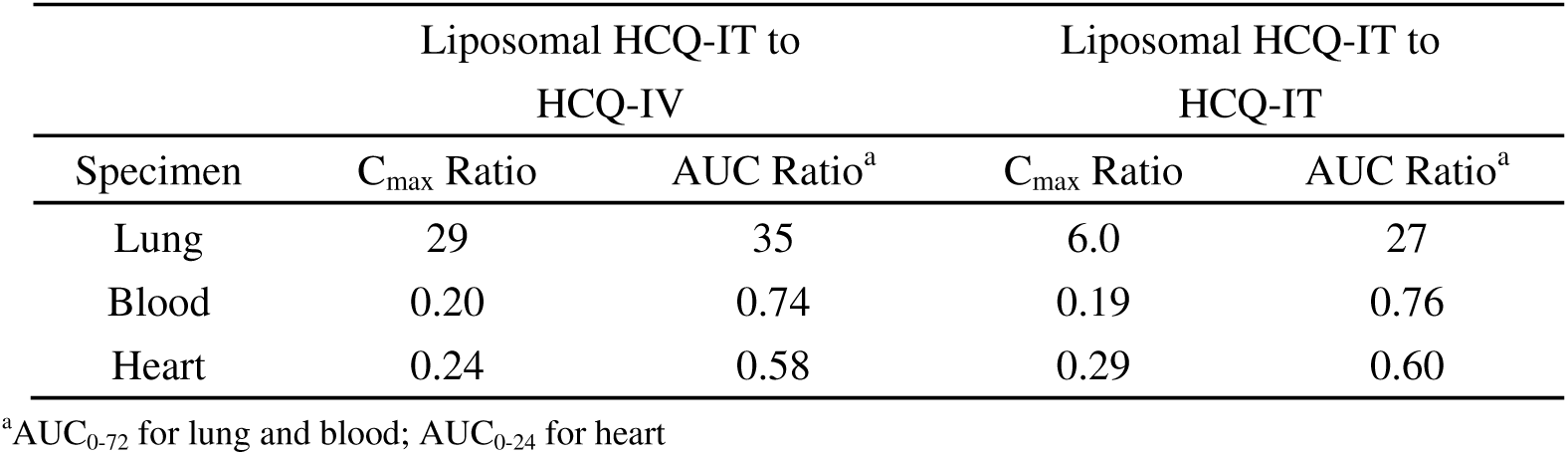
Ratios of dose-normalized maximum concentration and area under the concentration-time curve for liposomal HCQ-IT to HCQ-IV or to HCQ-IT.

**FIG 1.**
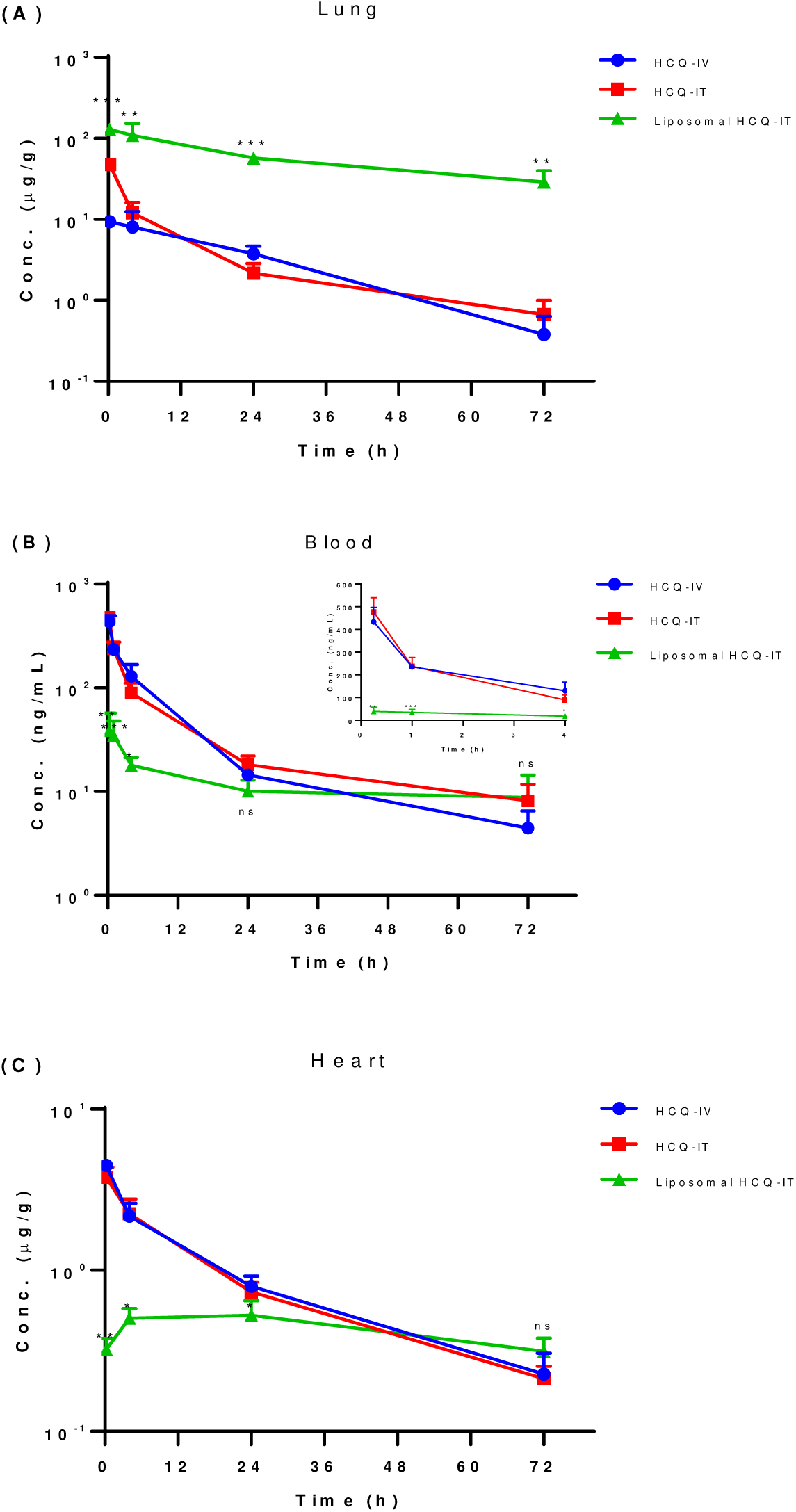
Mean concentration-time profiles of HCQ in rat lung (A), blood (B), and heart (C) after a single administration of liposomal HCQ through intratracheal delivery (IT) or HCQ through intravenous (IV) or IT. The inset graph (B) showed mean concentration-time profiles of HCQ in 0.25 to 4 hours. ^*^*P* < 0.05; ^**^*P* < 0.01; ^***^*P* < 0.001; ^ns^*P* > 0.05 compared to HCQ-IV.

### HCQ pharmacokinetics in blood and heart

With IV administration, HCQ showed similar PK profile and systemic exposures as HCQ-IT in overall, including C_max_ and AUC (Fig. 1B and Table 1). On the contrary, liposomal HCQ-IT showed significantly lower systemic exposure in blood with only around 20% of C_max_ and 74% of AUC_0-72_ after normalizing dose compared to HCQ-IV (Table 2). As observed in blood, liposomal HCQ-IT showed significantly lower exposure in the heart from 0.25 hours to 24 hours (Fig. 1C and Table 1). With normalized dose, liposomal HCQ-IT yielded only 24% of C_max_ and 58% of AUC_0-24_ compared to HCQ-IV (Table 2).

## DISCUSSION

In this rat pharmacokinetics study, a significantly higher exposure of HCQ with sustained release in the lung was observed by targeted delivery of inhalable liposomal HCQ, suggesting a potential treatment strategy for COVID-19 pulmonary disease. Fan et al. and others have suggested that significantly higher lung (intracellular) concentrations relative to the *in vitro* EC_50_ would be required to achieve *in vivo* antiviral efficacy SARS-CoV-2 (5, 6). Also, it has been proposed that the prediction of *in vivo* efficacy should be driven primarily by high lung HCQ concentrations for treatment of viral pneumonia instead of HCQ blood exposure (6). Given the current dosing regimen for oral HCQ is unlikely to produce an antiviral effect, FDA recently revoked the emergency use authorization (EUA) to use HCQ to treat COVID-19 in certain hospitalized patients and WHO also stopped the HCQ arm of the COVID-19 Solidarity Trial. Here our findings supported the feasibility of alternative targeted delivery of HCQ to the lungs with potentially effective antiviral levels while minimizing the systemic exposure.

The aerosolized delivery of therapeutic drugs to the lower respiratory tract has been applied for the treatment of various lung infectious and inflammatory disorders. For example, aerosolized HCQ (AHCQ) has been applied in clinical trials for asthma and showed well tolerance without significant toxicity after 21 days of dosing (8). In the current study, it is found that HCQ lung exposure remains low with short half-life even with dosing of unformulated HCQ directly to trachea, indicating unformulated HCQ is possibly to be retained in lung only transiently and distributed rapidly from lung to systemic circulation. The results suggest a sustained release formulation of HCQ, like inhalable liposomal HCQ demonstrated here might be able to provide preferentially higher concentrations in the lung than an aerosolized, non-liposomal HCQ formulation.

It was demonstrated that the inhalable liposomal formulation could provide efficient aerosolized delivery of liposomal drugs to lung and increase drug exposure in airways and lung with lower doses than used for systemic administration (9). For instance, ARIKAYCE^®^, an inhalable liposomal amikacin has been developed and accelerated approved by FDA in 2018 for treating Mycobacterium avium complex lung disease (10). As shown in this proof-of-concept study, the inhalable liposomal HCQ did increase exposure in the lung with extended residence time compared to systemically administered HCQ. Notably, the significantly increased lung exposure at normalized dose suggested the lung HCQ concentration with *in vivo* antiviral efficacy could be achieved at a relatively lower dose of inhalable liposomal HCQ than that of oral regimens with loading dose initiation (11).

In addition to prolonging the residence time and increasing the exposure in lung, significantly lower effective dose with less dosing frequency of inhalable liposomal HCQ might also reduce systemic exposure and potential adverse events. One of the well-known side effects of HCQ in the treatment of rheumatological disorders is cardiotoxicity, including abnormal heart rhythms such as QTc interval prolongation and ventricular tachycardia, a dangerously rapid heart rate (12). In recent study, among 84 COVID-19 patients who received HCQ and azithromycin at NYU’s Langone Medical Center, 11% of them had prolonged QT intervals to be considered at high risk of arrhythmia (13). The present study showed liposomal HCQ-IT not only reduced systemic exposure but also decreased heart tissue distribution of HCQ compared to HCQ-IV. It suggested inhalable liposomal HCQ at a significantly lower effective dose should possess less cardiotoxicity concerns than conventional oral HCQ tablets.

While there is no commercially available aerosolized formulation of HCQ, a recent empirical study of inhaled HCQ aerosols at 4 mg per day over one week found it was well tolerated without significant adverse events (14). This further supports the rationale of applying inhalable liposomal HCQ with direct lung targeting in COVID-19 infection/disease prevention and treatment.

A limitation of the current study is that the pharmacokinetics of liposomal HCQ were evaluated in a rat model using IT instillation to mimic the intended inhalation administration. The typical lung deposition efficiency with inhaled aerosols is very likely lower than that of the IT instilled microsprayed-droplets. Furthermore, it is suggested the aerosol-generating procedure (nebulization) on patients with known or suspected COVID-19 should be performed cautiously given that SARS-CoV-2 is highly contagious through the respiratory route (https://www.cdc.gov/coronavirus/2019-ncov/hcp/infection-control-recommendations.html) (15). Therefore, we have developed a disposable closed-loop system connected to the nebulizer to maximize the targeted delivery of inhaled liposomal HCQ while minimize the spreading and contaminating the air and environment (data not shown).

In conclusion, this study in a rat model demonstrate the desirable pharmacokinetics of inhalable liposomal HCQ *in vivo*. It supports the working hypothesis that inhalable liposomal HCQ might serve as a potential treatment option for the delivery of HCQ for COVID-19 pulmonary disease by achieving targeted antiviral levels with less frequent dosing and at a relatively lower dose.

## MATERIALS AND METHODS

### Drugs and reagents

The study drug (liposomal HCQ) was prepared by Taiwan Liposome Company, Ltd., Taiwan. It is composed of HCQ encapsulated in liposomes with mean particle size around 200 nm. The liposomes are composed of dipalmitoylphosphatidylcholine (Lipoid GMBH, Germany) and cholesterol (Carbogen Amcis B.V., The Netherlands), both of which are natural components of lung surfactant (16, 17). The formulation was manufactured in the following processes. Briefly, appropriate amounts of lipid mixture were dissolved in ethanol (J. T. Baker, USA) and injected to a hydroxychloroquine sulfate (SCI Pharmtech, Inc., Taiwan) solution at 50°C. The size of the liposomes was adjusted to 200 nm by an extruder with 0.2 µm polycarbonate membrane to standardize the sizes of the liposomes. A tangential flow filtration (TFF, VIVAFLOW 200, MWCO 100,000 PES, Sartorius Stedim Biotech GmbH, Germany) was used to remove non-incorporated HCQ and ethanol to obtain the liposomal HCQ in 0.9% sodium chloride (Merck KGaA, Germany).

### Study Design

A total of 52 female SD rats (BioLASCO Taiwan Co., Ltd.) were assigned to one of three treatment groups: (1) HCQ-IV: 12 rats received a single dose of 0.590 mg HCQ sulfate per animal via IV injection; (2) HCQ-IT: 20 rats received a single dose of 0.590 mg HCQ sulfate per animal via intratracheal (IT) administration; and (3) liposomal HCQ-IT: 20 rats received a single dose of 0.284 mg liposomal HCQ sulfate per animal via IT administration. The sampling time points for blood samples were 0.25, 1, 4, 24 and 72 hours post-dose and for tissue/organ samples were 0.25, 4, 24 and 72 hours post-dose. In this study, inhalable liposomal HCQ was administered through IT instillation to mimic the intended inhaled administration in a clinical setting. All procedures involving animals were performed in TLC animal facility and in accordance with the ethical guidelines of Institutional Animal Care and Use Committee (IACUC) at TLC, Taiwan (#TLC20IACUC012).

Blood was collected from jugular veins at scheduled sampling time points into collection tubes with K_2_EDTA as the anticoagulant and stored at -80°C until analysis. After blood draw, each animal was perfused with 2 mM K_2_EDTA/saline solution before lungs and hearts were removed, weighed and stored at -80°C.

### Bioanalysis and PK calculation

Blood samples were mixed well with acetonitrile with 0.1% formic acid for protein precipitation. The supernatant was dried by nitrogen and reconstituted with 30% acetonitrile with 0.1% formic acid containing internal standard (IS) prior to injection into a liquid chromatography-tandem mass spectrometer (LC-MS/MS). For lung and heart samples, tissue/organ was homogenized with 50% methanol with 0.1% formic acid. The tissue/organ homogenate was added with IS and mixed well with acetonitrile with 0.1% formic acid for protein precipitation. The resulting sample solution was mixed well with 0.1% formic acid to final 30% acetonitrile with 0.1% formic acid prior to injection into a LC-MS/MS.

Concentrations of HCQ in blood and tissue/organ samples were determined by LC-MS/MS on a Waters ACQUITY UPLC-I CALSS coupled with Water Xevo TQS or AB Sciex Triple Quad 5500. The ACQUITY BEH C18 column (2.1×50 mm) was selected for sample separation with gradient elution consist of 0.1% formic acid in water (mobile phase A) and acetonitrile containing 0.1% formic acid (mobile phase B) at 0.4 mL/min flow rate. Total run time was 4.5 minutes and the column oven was set at 40°C. The mass spectrometer was set up in MRM mode to monitor the transition 336 → 247 for HCQ. The linear range of the assay is 0.5-500 ng/mL, 0.5-500 ng/mL and 20-10,000 ng/mL for blood, heart and lung assay, respectively. PK parameters of HCQ were calculated by a non-compartmental method using Phoenix^®^ WinNonlin^®^ (version 8.0). Sparse sampling computation was applied for the calculation of PK parameters in tissue/organ.

### Statistical Analysis

Two-sample *t* tests were used to assess differences in concentration between liposomal HCQ-IT and HCQ-IV. *P* < 0.05 is considered significant.

## ACKNOWLEDGEMENTS

We thank Dr. George Spencer-Green for his critical review and suggestions for the manuscript. We also thank Ting-Yu Cheng for the formulation development, staff in Department of Pharmacokinetics, Taiwan Liposome Company for the laboratory support, and staff in Department of Nonclinical, Taiwan Liposome Company for the support of animal model. This study was supported by Taiwan Liposome Company, Ltd., Taiwan.

